# (Epi-)Genomic Data in the German TwinLife Study: TwinSNPs and TECS Cohort Profiles

**DOI:** 10.64898/2026.02.20.704007

**Authors:** Leonard Frach, Charlotte K. L. Disselkamp, Alicia M. Schowe, Anastasia Andreas, Marco Deppe, Jana Instinske, Carlo Maj, Theresa Rohm, Mirko Ruks, Lucia M. Wiesmann, Christian Kandler, Bastian Mönkediek, Frank M. Spinath, Markus M. Nöthen, Elisabeth B. Binder, Darina Czamara, Andreas J. Forstner

**Author notes:** Correspondence &. shared first authors (Leonard Frach, Charlotte K. L. Disselkamp, Alicia M. Schowe). shared senior authors (Darina Czamara, Andreas J. Forstner).

## Abstract

The German Twin Family Panel *TwinLife* is a nationwide longitudinal study of twins and their family members. Primarily focusing on the development of social inequalities over the life course, TwinLife has been collecting data since October 2014 starting with 4,096 twin families (*N_total_* = 16,951 individuals). As Germany’s largest twin study to date, TwinLife has been surveying four birth cohorts of monozygotic and dizygotic same-sex twin pairs (initially ∼5, 11, 17, and 23 years old) and their families for 11 years. Survey data have been collected through five biennial face-to-face interviews with four computer-assisted telephone interviews in the years between. In addition, saliva samples were collected before the COVID-19 pandemic (2018-2020), during the pandemic (2021), and after (2022-2024). In this Cohort Profile, we describe the curation and initial analyses of molecular genetic and epigenetic data from the two TwinLife satellite projects TwinSNPs and TECS. Together, these projects currently comprise 12,108 processed DNA samples from 6,450 participants, extracted from the first two saliva collections before and during the COVID-19 pandemic. We compared the subsamples with the overall TwinLife sample and provide an overview of derived polygenic scores (PGS), epigenetic clocks and other methylation profile scores (MPS). We found that PGS predicted sample attrition in TwinLife, with small but significant associations between higher PGS for educational attainment and continued participation. Epigenetic clocks derived from saliva were highly correlated with chronological age (*r* = .71 to *r* = .94) and were generally more stable over time than other MPS. PGS for epigenetic clocks were associated with the respective clock only during but not before the start of the pandemic. We discuss opportunities of combining prospectively assessed molecular (epi)genetic data in within-family designs such as TwinLife and its implications and avenues for future research.

## Introduction

*TwinLife* was established in 2014 at the Bielefeld and Saarland University, with the University of Bremen joining the project in 2021. It is the first twin family study in Germany to employ a population register-based sampling design (1,2). The TwinLife study was set up to investigate the development of social inequalities in a genetically informative design. To complement its rich longitudinal family-based design with biological sampling, the prospective collection of saliva samples was integrated and led to the TwinSNPs satellite project. TwinSNPs aims to combine quantitative genetic methods with molecular genetic data. The addition of molecular genetic data to an extended twin family study enables further insights into genetic and environmental contributors to social inequality and related outcomes. The acute societal challenges of the COVID-19 pandemic prompted the launch of the TwinLife Epigenetic Change Satellite (TECS) project in 2021, which includes a sub-sample of genotyped twins from the TwinSNPs sample. The TECS project investigates how pandemic-associated stressors affect the epigenome, specifically DNA methylation (DNAm), and, in turn, physical, mental, and social health. It focuses on impactful stressors, moderators of epigenetic changes, and risk versus resilience profiles by further integrating genetic, psychological, and social factors.

This Cohort Profile describes the integration, curation, and initial analyses of genetic and epigenetic data collected in the German Twin Family Panel TwinLife through the TwinSNPs and TECS satellite projects. The aim is to provide an overview of available genotyped samples, polygenic scores (PGS), epigenetic clocks and other methylation profile scores (MPS), enabling future research on genetic, epigenetic, and environmental influences on development and social inequality across the life course.

## Sample Description

TwinLife started data collection in October 2014, including same-sex twin pairs reared together and born in 2009/2010 (cohort 1), 2003/2004 (cohort 2), 1997/1998 (cohort 3), and between 1990 and 1993 (cohort 4). It used a register-based probability sampling design including families from across Germany, including communities with ≥ 5,000 inhabitants (1) and families from diverse socioeconomic backgrounds (3). In addition to the twin pairs, data have been collected on their parents, the parents’ partners (e.g., non-biological parents) and non-twin siblings, enabling an extended twin family design (Supplementary Figure S1) (4). In addition, for adult twins, there is phenotypic data available on the partners and children of twins. Molecular genetic data of the TwinSNPs sample have been collected for twins, siblings, and biological parents, whereas DNAm data in the TECS sample is currently only available for twins who participated in the COVID-19 pandemic supplementary surveys (Cov), who had already been genotyped and had passed quality control of the genetic data to maximise the overlap of molecular data (see below for details). An overview of the current state of the TwinLife study, phenotypic measures and other satellite projects beyond TwinSNPs and TECS can be found in Mönkediek et al. (5).

## Data Collection and Measures

Phenotypic data were collected in face-to-face interviews (F2F), computer-assisted telephone interviews (CATI), and additional surveys during the pandemic (Cov) as shown in Figure 1. TwinLife covers an extensive range of phenotypes. In addition to socioeconomic information, phenotype data include psychological measures on personality, subjective well-being, externalising and internalising problems and intelligence tests, among others (1,6,7). See Mönkediek et al. (5) for the most up-to-date overview of the project and the phenotypic data, which can also be found online: https://www.twin-life.de/documentation/. Additional information on specific scales in TwinLife can be found in the scales manual (8).

**Figure 1.**
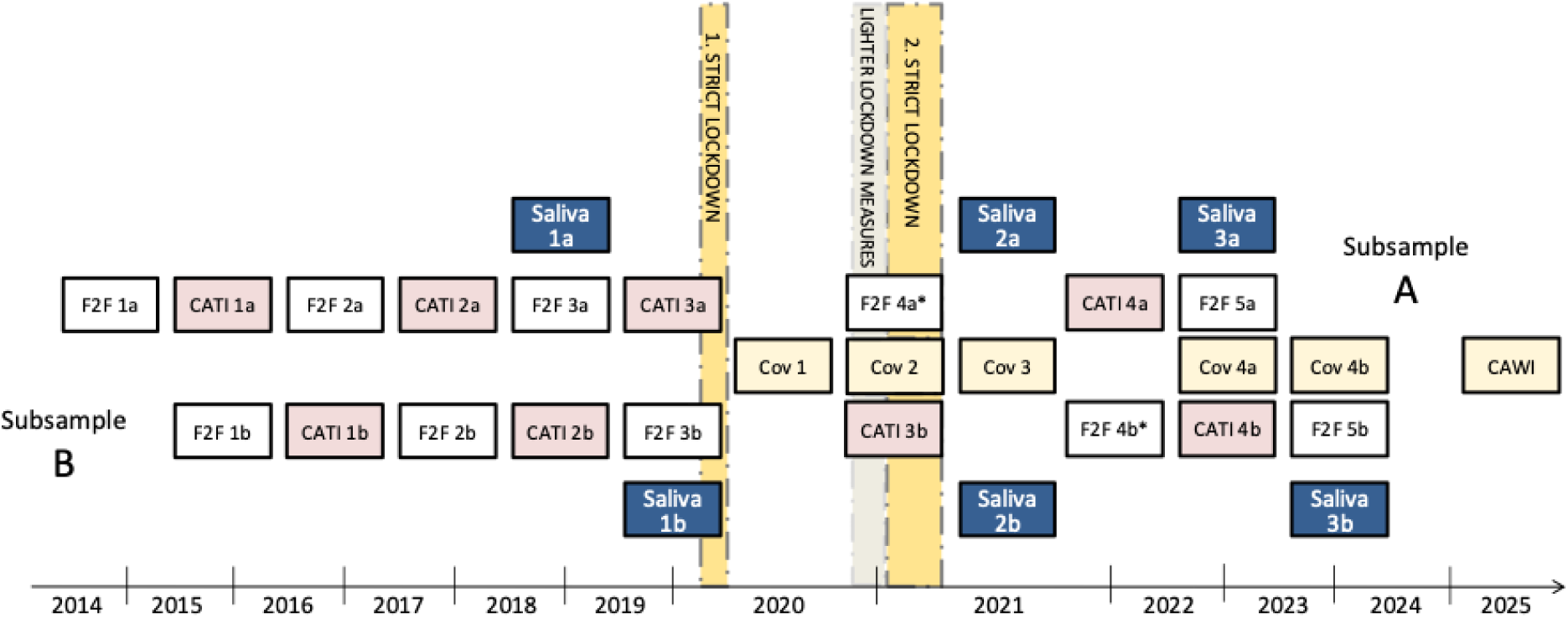
Data Collection Timepoints of Saliva Samples and Phenotypic Data. *Note.* Data collection waves, including face-to-face interviews (F2F), computer-assisted telephone interviews (CATI), and COVID-19 pandemic supplementary surveys (Cov). Collection was divided into two subsamples, with data collection of subsamples A and B alternated each year. Saliva = collection of saliva via self-collection kit. Figure adapted from Mönkediek et al. (5). *Fourth face-to-face (F2F4) interview was replaced by a CATI & CAWI (computer assisted web interview) due to the COVID-19 pandemic. F2F4 was also for re-assessment of the first saliva sample of individuals who consented to the saliva sampling but without available saliva sample from F2F3.

### Saliva Sampling and Processing

At F2F3 and Cov3, the first two saliva samples of participants were collected using self-collection kits. For individuals who consented to provide a saliva sample but did not have an available sample for F2F3, the first saliva sample was assessed at F2F4. Additional saliva samples were collected as part of F2F5 (see (5) for more details). Processing of additional saliva samples from F2F5 is ongoing. The collection of saliva samples was announced to participants in the letters inviting them to participate in the survey waves. Respondents were provided with informed consent forms, Oragene® saliva self-collection kits (OG-500 or OG-600; DNA Genotek, Canada), and instructions. At F2F3 the interviewers introduced the TwinSNPs satellite project to participants, provided them with information materials and showed them a short information video including instructions on how to collect the saliva samples. All materials were also made available on the study website, the link to which was provided in the cover letter, so that participants could obtain information in advance. At later waves, twins and their family members received further invitations and self-collection kits by mail.

DNA extraction, genotyping and further genomic data processing were performed at the Institute of Human Genetics and the Institute for Genomic Statistics and Bioinformatics of the University of Bonn and the University Hospital Bonn, Germany. All individuals that consented to provide DNA from saliva samples during any of the timepoints F2F3, F2F4 or Cov3 are part of the current TwinSNPs cohort. For its molecular genetic satellite projects TwinSNPs and TECS, a total of 12,742 samples from 6,542 individuals have been collected (including twins and their family members, including timepoint F2F5), of which 12,108 saliva samples from 6,450 individuals have been processed to date. For the TECS sample, DNAm was analysed at F2F3 and Cov3 waves, as shown in Figure 1.

### Sample Attrition

TwinLife started with 4,096 twin families during the first interview. As described in previous publications in detail (1,5,6,9), over the course of the study, TwinLife experienced selective attrition in terms of educational and migration background (6), which is commonly observed in longitudinal studies. However, the decline was exacerbated by a necessary change of survey institute during the second wave of the survey. This led to families who continued study participation being more highly educated and having a higher household income (Table 1). This is also reflected in the characteristics of the participants of the two satellite projects TwinSNPs and TECS. Compared to F2F1 of the TwinLife study, an even larger proportion of highly educated participants took part in the satellite projects described here (Table 1). In addition, individuals predominantly had no migration background (Table 1). The latter is partly because there was a lower share of individuals with migration background among those who contributed a saliva sample compared to F2F1 (Table 1). Furthermore, due to the removal of population outliers during the genetic quality control (individuals with ± 6 SD away from the mean on the first 10 principal components, PCs), the TwinSNPs sample is currently restricted to individuals of European ancestry (according to the 1000 Genomes reference sample). Notably, sampling weights have been created for TwinLife to account for unequal sampling probabilities due to the sampling design, and potential nonresponse bias due to selective nonresponse in F2F1 and selective attrition in the subsequent waves (10). Further analyses and results of sample representativeness and attrition using genetic data are shown in the section ‘Polygenic Scores Predict Sample Attrition’ below.

**Table 1.** Sample Characteristics of Twins from the Whole TwinLife Sample at Baseline (F2F1) and the TwinSNPs and TECS Subsamples.

| **Variable** | **F2F1** | **Valid Saliva Sample** | **TwinSNPs all<sup>a</sup>** | **TwinSNPs complete<sup>b</sup>** | **TECS all<sup>a</sup>** | **TECS complete<sup>b</sup>** |
| --- | --- | --- | --- | --- | --- | --- |
| ***N* Twins** | 8,192 | 3,304 | 2,718 | 2,426 | 1,055 | 994 |
| ***N* Families** | 4,096 | 2,262 | 1,619 | 1,213 | 558 | 497 |
| **Sex** |  |  |  |  |  |  |
| *Female* | 4,484 (55%) | 1,809 (55%) | 1,467 (54%) | 1,314 (54%) | 561 (53%) | 524 (53%) |
| *Male* | 3,708 (45%) | 1,495 (45%) | 1,251 (46%) | 1,112 (46%) | 494 (47%) | 470 (47%) |
| **Migration Background<sup>c</sup>** |  |  |  |  |  |  |
| *No* | 3,430 (76%) | 1,596 (80%) | 1,414 (84%) | 1,280 (84%) | 648 (85%) | 607 (85%) |
| *Yes* | 1,058 (24%) | 399 (20%) | 264 (16%) | 247 (16%) | 114 (15%) | 108 (15%) |
| **Age in F2F1** | 14.0 (6.7) | 13.1 (6.6) | 13.1 (6.6) | 12.7 (6.4) | 11.4 (6.1) | 11.4 (6.0) |
| **Birth Cohort** |  |  |  |  |  |  |
| *Cohort 1* | 2,018 (25%) | 928 (28%) | 763 (28%) | 724 (30%) | 366 (35%) | 338 (34%) |
| *Cohort 2* | 2,086 (25%) | 926 (28%) | 775 (29%) | 728 (30%) | 395 (37%) | 376 (38%) |
| *Cohort 3* | 2,122 (26%) | 801 (24%) | 643 (24%) | 558 (23%) | 161 (15%) | 154 (15%) |
| *Cohort 4* | 1,966 (24%) | 649 (20%) | 537 (20%) | 416 (17%) | 133 (13%) | 126 (13%) |
| **Zygosity** |  |  |  |  |  |  |
| *DZ* | 4,364 (53%) | 1,688 (51%) | 1,390 (51%) | 1,222 (50%) | 493 (47%) | 462 (46%) |
| *MZ* | 3,822 (47%) | 1,616 (49%) | 1,328 (49%) | 1,204 (50%) | 562 (53%) | 532 (54%) |
| **Smoking Behaviour in F2F1<sup>d</sup>** |  |  |  |  |  |  |
| *(Heavy) Smoker* | 514 (13%) | 141 (9.8%) | 121 (10%) | 96 (9.9%) | 16 (5.5%) | 16 (5.8%) |
| *Light Smoker* | 179 (4.4%) | 65 (4.5%) | 51 (4.3%) | 41 (4.2%) | 7 (2.4%) | 7 (2.5%) |
| *Former Smoker* | 477 (12%) | 148 (10%) | 130 (11%) | 105 (11%) | 26 (8.9%) | 23 (8.3%) |
| *Never Smoked* | 2,880 (71%) | 1,087 (75%) | 871 (74%) | 726 (75%) | 243 (83%) | 232 (83%) |
| **BMI in F2F1<sup>3</sup>** | 19.3 (4.5) | 18.8 (4.4) | 18.8 (4.5) | 18.5 (4.4) | 17.7 (4.2) | 17.8 (4.3) |

**Table 1 continued**
| **Variable** | **F2F1** | **Valid Saliva Sample** | **TwinSNPs all<sup>a</sup>** | **TwinSNPs complete<sup>b</sup>** | **TECS all<sup>a</sup>** | **TECS complete<sup>b</sup>** |
| --- | --- | --- | --- | --- | --- | --- |
| **Community Size in F2F1** |  |  |  |  |  |  |
| *5,000-19,999 inhabitants* | 1,543 (19%) | 638 (19%) | 548 (20%) | 496 (20%) | 221 (21%) | 213 (21%) |
| *20,000-99,999 inhabitants* | 1,829 (22%) | 740 (22%) | 594 (22%) | 520 (21%) | 234 (22%) | 216 (22%) |
| *100,000-499,999 inhabitants* | 2,291 (28%) | 939 (28%) | 776 (29%) | 700 (29%) | 334 (32%) | 317 (32%) |
| *500,000+ inhabitants* | 2,525 (31%) | 986 (30%) | 800 (29%) | 710 (29%) | 266 (25%) | 248 (25%) |
| **Parental Education (ISCED) in F2F1** |  |  |  |  |  |  |
| *ISCED 1* | 120 (1.5%) | 24 (0.7%) | 10 (0.4%) | 8 (0.3%) | 0 (0%) | 0 (0%) |
| *ISCED 2* | 424 (5.2%) | 70 (2.1%) | 30 (1.1%) | 24 (1.0%) | 2 (0.2%) | 0 (0%) |
| *ISCED 3* | 2,210 (27%) | 696 (21%) | 554 (20%) | 480 (20%) | 179 (17%) | 172 (17%) |
| *ISCED 4* | 542 (6.7%) | 223 (6.8%) | 197 (7.3%) | 176 (7.3%) | 70 (6.6%) | 62 (6.2%) |
| *ISCED 5a* | 2,536 (31%) | 1,179 (36%) | 988 (36%) | 898 (37%) | 404 (38%) | 380 (38%) |
| *ISCED 5b* | 1,846 (23%) | 886 (27%) | 764 (28%) | 682 (28%) | 309 (29%) | 292 (29%) |
| *ISCED 6* | 462 (5.7%) | 217 (6.6%) | 171 (6.3%) | 156 (6.4%) | 91 (8.6%) | 88 (8.9%) |
| **OECD Household Net Income in F2F1** | 1,658 (1,265) | 1,780 (1,295) | 1,822 (1,353) | 1,836 (1,368) | 1,972 (1,548) | 1,987 (1,580) |

*Note.* Numbers show the *N* with % in parentheses, or the mean values and standard deviation in parentheses (for age and income). F2F = face-to-face interview, ISCED = International Standard Classification of Education.
<sup>a</sup> including complete twin pairs and individual twins, as well as additional family members (TwinSNPs only).
<sup>b</sup> complete twin pairs, i.e., data is available for both twins.
<sup>c</sup> Following the definition of the German federal statistical office, a person has a migration background if they or at least one parent was not born with German citizenship. Thus, migration background is only calculated for persons with valid information on citizenship for both themselves and their parents.
<sup>d</sup> Smoking behaviour and BMI (body mass index) have not been assessed at F2F3 (the timepoint of the first DNAm assessment) but baseline information can be used as a proxy for later timepoints. Note that smoking was not assessed for all twins at baseline due to the young age of cohorts 1 and 2.

**Table 2.** Composition of Family Members Compared Between TwinLife Sample at Wave 1 and TwinSNPs and TECS Subsamples After QC.

|  | **TwinLife F2F1** | **TwinSNPs** | **TECS<sup>a</sup>** |
| --- | --- | --- | --- |
| *N* complete twin pairs | 4,096 | 1,213 | 497 |
| *N* additional twins | – | 292 | 69 |
| *N* non-twin siblings | 2,106 | 522 | – |
| *N* fathers | 3,667 | 917 | – |
| *N* mothers | 4,051 | 1,263 | – |

<sup>a</sup> Epigenetic data are currently only available for twins. Genetic and phenotypic data are available for parents and siblings of TECS members, in addition to twins.

### Genotyping and Quality Control

Genomic DNA was extracted from saliva samples for genotyping and assessment of DNAm according to the Oragene® saliva self-collection kits (OG-500 and OG-600; DNA Genotek, Canada) protocol. Genotyping was conducted using Illumina Global Screening Arrays with additional marker content relevant for psychiatric disorders (GSA+MD-24v3.0-Psych-24v1.1, Illumina, San Diego, CA, USA). Standard quality control (QC) of genetic data was performed following existing pipelines for family based samples (11) using PLINK v1.9 (12). In brief, individuals with sex mismatches, genotyping rate < 0.98 and heterozygosity rate >0.20 were removed. Variants with a call rate < 0.98, deviating from Hardy-Weinberg equilibrium with a *p*-value < 1 × 10^-6^ or with a minor allele frequency < 0.001 were initially removed. KING software (13) was used to confirm the known family structure of the TwinLife samples. Genotypes were phased using Eagle v2.4.1 (14) and imputed using Minimac v4 (15), with 1000 Genomes project phase 3 (1000Gv3) data as imputation reference panel (16). 1000Gv3 was also used for projecting TwinSNPs individuals onto the PC space using the first 10 PCs to infer genetic ancestry. See the Supplementary Note for more details. After QC, genome-wide genotype data were available for *N* = 5,421 TwinSNPs participants (see Table 3 for the family structure of genotyped individuals).

**Table 3.** Overview of genotyped twin-pairs and parent-offspring constellations.

|  | ***N Families*** |
| --- | --- |
| Complete MZ twin pairs | 570 |
| Complete DZ twin pairs | 643 |
| Independent trios | 788 |
| Independent mother-child duos | 1,196 |
| Independent father-child duos | 872 |
| Spouses | 806 |

### Polygenic Scores

Following additional QC of genome-wide association studies (GWAS) summary statistics (Supplementary Note), we computed PGS using PRS-CS (17) with default parameters (PRS-CS auto) and the 1000Gv3 European LD reference panel (see Supplementary Note for details). For analyses presented in this Cohort Profile, we used PGS for educational attainment (18), intelligence (18), depression (19) as well as height (20) as negative control (see sections ‘Polygenic scores predict sample attrition’ and ‘Selective sampling of TwinSNPs and TECS subsamples is reflected in polygenic scores’). Furthermore, we also present analyses using PGS for the epigenetic clocks Horvath, Hannum and GrimAge, derived from existing GWAS summary statistics (21), see section ‘Associations between polygenic scores and epigenetic clocks’. In addition, we calculated ready-to-use PGS covering various phenotypes ranging from educational and cognitive traits to personality as well as mental and physical health indicators based on publicly available GWAS summary statistics. For an overview and full list of currently available PGS in TwinLife see Table 4 and Supplementary Table S2 as well as the TwinLife website https://www.twin-life.de/documentation/generated-variables-and-scales. Furthermore, additional measurement error-corrected PGS (accounting for imperfect reliability of PGS (22)) are provided as part of the PGI repository v2 (23).

**Table 4.** Overview of Currently Available Polygenic Scores for TwinSNPs.

| Category | Polygenic Scores |
| --- | --- |
| Educational & cognitive traits | Educational attainment (including cognitive and non-cognitive aspects), cognitive performance, intelligence, reading & language abilities (word reading, IQ, spelling, non-word reading, non-word repetition, phoneme awareness) |
| Psychiatric disorders | ADHD, schizophrenia, bipolar disorder (including type 1 & 2), major depressive disorder, depressive symptoms, anxiety, post-traumatic stress disorder, panic disorder, Tourette syndrome, autism spectrum disorder, anorexia nervosa, alcohol dependence, alcohol use disorder, cannabis use disorder, opioid dependence, cross-disorder, obsessive-compulsive disorder |
| Well-being & personality traits | Subjective well-being, life satisfaction, positive affect, well-being spectrum, eudaimonic & hedonic well-being, extraversion, neuroticism, conscientiousness, agreeableness, openness, risk-taking, loneliness, financial satisfaction, family satisfaction, friendship satisfaction, work satisfaction |
| Physical traits | Height, body mass index, chronotype, morning person, epigenetic clocks (Hannum, Horvath, GrimAge, PhenoAge) |
| Inflammatory & biological markers | C-reactive protein, interleukin-10, interleukin-6, tumour necrosis factor-alpha, PAI-1 & granulocyte proportions |
| Other | Childhood maltreatment, self-rated health |

*Note.* See Supplementary Table S2 for an exhaustive list including references.

### Quality Control and Pre-processing of DNA Methylation Data

DNAm was measured using the DNA of 2,128 samples (1,064 twins with two samples each). Samples were randomised by age, sex, zygosity, and complete versus single twin pair across 23 plates (266 arrays) using the R package *Omixer* v1.4.0 (24). To prevent systematic batch effects within twin pairs and across repeated measures, all samples from each twin pair were grouped by family ID and processed on the same array. Assignment of families to arrays and individual sample positions within arrays were randomised. Bisulfite conversion of 500 ng DNA was performed using the D5033 EZ-96 DNA Methylation-Lightning Kit (Deep-Well; Zymo Research Corp., Irvine, USA) and methylation profiles were assessed using the Infinium MethylationEPIC BeadChip v1.0 (Illumina, San Diego, CA, USA). Preprocessing of all methylation samples (*N* = 2,128) was conducted following the steps by Maksimovic et al. (25,26). Specifically, scan intensity signals as stored in .idat files were loaded into R and transformed into beta values within the R package *minfi* v1.40.0 (27). Samples with a mean detection *p*-value > 0.05 were excluded (*N* = 3). Additionally, we excluded samples showing sex mismatches between estimated sex from methylation data and genotype-confirmed sex (*N* = 14). Beta values were normalised using stratified quantile normalisation (R package *minfi*, v1.40), followed by beta-mixture quantile (BMIQ) normalisation (R function published by Teschendorff et al. (28) v1.6). Probes on sex chromosomes (*n* = 16,263), probes containing SNPs (*n* = 24,038, annotated using R package *minfi*), and probes with a detection *p*-value > 0.01 in one or more samples (*n* = 79,922) were removed. We also excluded cross-hybridising (*n* = 46,867) and polymorphic probes (*n* = 297) identified by Chen et al. (29) and McCartney et al. (30), as well as probes within 10 base pairs of common SNPs (MAF > 0.05) noted in the Illumina Epic v1 manifest file (*n* = 36,160). Afterwards, beta-values were transformed into M values, and batch effects were removed using *ComBat* as implemented in the R package *sva* v3.46 (31). For this, we computed a principal component analysis (PCA) on the M values and checked which batch variables (plate, slide, array) were most strongly associated with the first five principal components. The strongest batch effects (plate and position) were iteratively removed. Corrected M values were re-transformed into beta values. Lastly, we applied *MixupMapper* v1.2.4 (32) to the genotype and methylation data to check for possible sample mix-ups. Four mix-ups were detected. For these four cases, both samples (baseline and follow-up) were removed (total of *N* = 8 individual samples). The final TECS sample consisted of *N* = 2,102 samples of *N* = 1,055 participants (of which *N* = 489 were complete twin pairs, 250 MZ and 239 DZ pairs with DNAm data at two timepoints) and *n* = 662,312 probes. Note that previous publications further included adjustment for array (33–35), which is no longer considered due to oftentimes unintuitive negative within twin-pair correlations at the CpG level. However, discrepancies between both approaches at the level of composite epigenetic scores were negligible.

### Epigenetic Clocks and Other Methylation Profile Scores

MPS are defined as the weighted sums of an individual’s methylation markers’ beta values of a pre-selected number of CpG sites. MPS can, for example, be used as biomarkers of aging, biomarkers of environmental exposures such as smoking, or to predict diseases or treatment success. We used the normalized and batch-corrected beta values without any probe filtering to compute 16 MPS and to estimate the salivary cell-type composition (see Table 5, Supplementary Table S2 and the TwinLife website https://www.twin-life.de/documentation/generated-variables-and-scales).

**Table 5.** Overview of Currently Available Methylation Profile Scores (MPS) for TECS.

| Category | MPS |
| --- | --- |
| Educational & cognitive traits | Epigenetic-g |
| Physical traits | Body mass index, puberty |
| Protein biomarkers | C-reactive protein |
| Lifestyle exposure | Smoking CpG methylation |
| Aging: biological markers | Telomere length, GrimAge, PC GrimAge, GrimAge2, PhenoAge, PC phenoAge, DunedinPACE |
| Aging: chronological age | Horvath DNAmAge, PC Horvath, SkinHorvath, PC SkinHorvath, BLUP, EN, PedBE, Wu, Hannum, PC Hannum |
| Other | Zygosity |

*Note.* BLUP = best linear unbiased prediction, EN = elastic net, PedBE = Pediatric-Buccal-Epigenetic

Cell-type composition was estimated using the R package *EpiDish* v2.10 (36), resulting in three estimated cell-type proportions: immune cells (T1: *M* = 0.75, *SD* = 0.09; *M* = 0.71, *SD* = 0.12), epithelial cells (T1: *M* = 0.23, *SD* = 0.09; T2: *M* = 0.28, *SD* = 0.12), and fibroblasts (T1: *M* = 0.02, *SD* = 0.01; T2: *M* = 0.02, *SD* = 0.01). Since these proportions are highly correlated, we also summarized cell-type variation in PCs which can be included as covariates in downstream analyses. The first PC explained 99.5 and 99.7% of the variation in cell-type composition at timepoint 1 and timepoint 2, respectively.

MPS were computed based on the prediction model provided by the respective original publication or as the weighted sum of beta values (see Supplementary Table S1). To date, calculated MPS include 10 chronological epigenetic clocks, seven biological epigenetic clocks, two MPS for protein biomarkers (i.e., CRP), two MPS for physical traits (BMI, puberty), one MPS for general cognitive ability, and one MPS for DNAm-based prediction of zygosity. For epigenetic age estimates, we identified one sample as an outlier (Horvath DNAmAge and PedBE +4 *SD* above the cohort mean) which was thus excluded from the epigenetic clock data prior to performance evaluation and any subsequent analyses. We evaluated MPS performance in the full TECS sample (*N* = 1,055 for MPS, *N* = 1,054 for epigenetic clocks) based on their correlation with the corresponding phenotype, when available (see Supplementary Table S1 and Supplementary Note). Epigenetic clocks were rather stable over time with intra-class coefficients (ICC) ranging between 0.36 to 0.95 and highly correlated with chronological age (Pearson’s *r* ranging between 0.71 and 0.94). The least stable clock was DunedinPACE which may indicate greater sensitivity to environmental influences. In contrast to all the other clocks, DunedinPACE was trained on longitudinal biomarkers of aging and represents pace of aging. All other epigenetic clocks were trained on cross-sectionally collected biomarkers or chronological age and represent epigenetic age (in years). For these latter clocks, we also provide the median absolute error (MAE, i.e., the absolute median difference between chronological and epigenetic age) and median deviation (i.e., the median between chronological and epigenetic age) separately for each age group to examine potential cohort-specific biases in epigenetic age estimation (see Supplementary Figures S7-S8). ICCs of MPS ranged between 0.32 and 0.90, with lowest stability for the MPS for CRP and the highest stability for the MPS for zygosity. Notably, predictive performance of the zygosity was similar in our saliva sample (82.26% and 83.33% correctly identified MZ twins at T1 and T2) compared to the performance in blood in the original study (84.30% correctly identified MZ twins).

## Recent Findings Using TwinSNPs and TECS Data

A comprehensive list of publications using the TwinLife data is provided at https://www.twin-life.de/publikationen. Using molecular genetic data from the TwinSNPs sample, first work showed significant gene-environment interactions for life satisfaction. That is, the PGS for subjective well-being showed the strongest association with life satisfaction in individuals from moderately deprived regions (37). Other work showed significant associations between PGS for personality traits and subjective well-being of twins, and between PGS and corresponding phenotypic measures of personality and well-being (38). In addition, partner similarity among twin-parents’ educational levels provided evidence for assortative mating and was partly explained by similarity on education-related PGS (39). To date, the TwinSNPs sample has already contributed to two published and several ongoing multi-cohort studies and meta-analyses, e.g., within-family PGS analyses (23,40,41). Ongoing efforts include, for example, a within-family trio GWAS, a GWAS of dizygotic twinning (https://www.twinningconsortium.org/members/) or a trio PGS meta-analysis of offspring externalising and internalising behaviours.

Analyses of epigenetic data from the TECS sample revealed that most variation in biological aging (measured by epigenetic clocks derived from DNAm) was explained by non-shared environmental factors alongside significant contributions of genetic and shared environmental effects (33). In addition, a longitudinal analysis using TECS data found that several epigenetic clocks (PedBE and Horvath epigenetic age acceleration and Dunedin pace of aging) increased from the first to the second saliva sampling timepoint. The increase in pace of aging was linked to higher perceived pandemic burden only in adolescents, suggesting that mid to late adolescence may be a sensitive period for subjective stress-related alterations in biological aging (34). Furthermore, recent research showed that acceleration in epigenetic aging is linked to change in personality, which is explained by a dynamic interplay of genetic influences on baseline differences and environmental influences on individual change (35). TECS data were also used in a multi-cohort study, confirming the validity of newly developed MPS for pubertal development to be associated with earlier age at menarche (40). Two ongoing multi-cohort efforts include association analyses between a PGS and MPS for cognitive ability (developed in adults) and cognitive and academic performance in children as well as a validation study of a newly developed MPS for poverty and socioeconomic status. Ongoing epigenome-wide analyses examine genetic and environmental influences on salivary DNAm at individual CpG sites, their stability over time, and the identification of methylation quantitative trait loci (mQTLs)—representing the first application of both classical twin modelling and QTL analysis to saliva-based DNAm data (https://osf.io/7g4aq).

## Polygenic Scores Predict Sample Attrition

To further assess selective attrition using genotyped individuals from TwinSNPs, we examined associations between PGS for educational attainment (EA), intelligence and depression, and height (as negative control) and study participation in F2F4 (drop-out N = 81 twins) and F2F5 (drop-out N = 109 twins). For all analyses including related individuals, we calculated clustered standard errors using the family ID. Using logistic regressions adjusting for genetic PCs, sex, and cohort, we found that PGS for EA and intelligence significantly predict study drop-out, i.e., individuals who continued participation had higher PGS for EA and intelligence compared to individuals who did not participate at F2F4. Specifically, we found that PGS for EA (OR = 1.34, 95% CI = [1.04, 1.73]) significantly predicted participation of twins at F2F4 (see Figure 2). For all individuals (twins, siblings and parents) both PGS for EA (OR = 1.31, 95% CI = [1.13, 1.52]) and intelligence (OR = 1.17, 95% CI = [1.01, 1.36]) predicted continued participation, but PGS for depression and height did not (OR = 1.00, 95% CI = [0.86, 1.16] and OR = 1.11, 95% CI = [0.95, 1.29], respectively). This indicates that individuals who remained in the study have higher PGS for EA and intelligence, which mirrors the higher observed education of continuing participants at the phenotypic level ((42), see also Table 1). Such a pattern has also been shown in other longitudinal cohort studies using PGS (43). Albeit showing similar trends, none of the PGS significantly predicted participation at F2F5 (see Figure 2), which may be due to the different absolute numbers in participation and thus resulting in lower statistical power. Interestingly, F2F4 data were assessed using telephone and computer assisted web interviews instead of in-person due to the COVID-19 pandemic, which is however similar to previous CATI interviews (Figure 1). Our results may indicate that selective attrition related to mental health and socioeconomic factors was amplified during the pandemic, which might be due to an increased pandemic-related burden for individuals of already disadvantaged groups. In addition to existing sampling weights, this information can be used for multiple imputation of phenotype data at follow-up timepoints, e.g., by including PGS as auxiliary variables (44,45).

**Figure 2.**
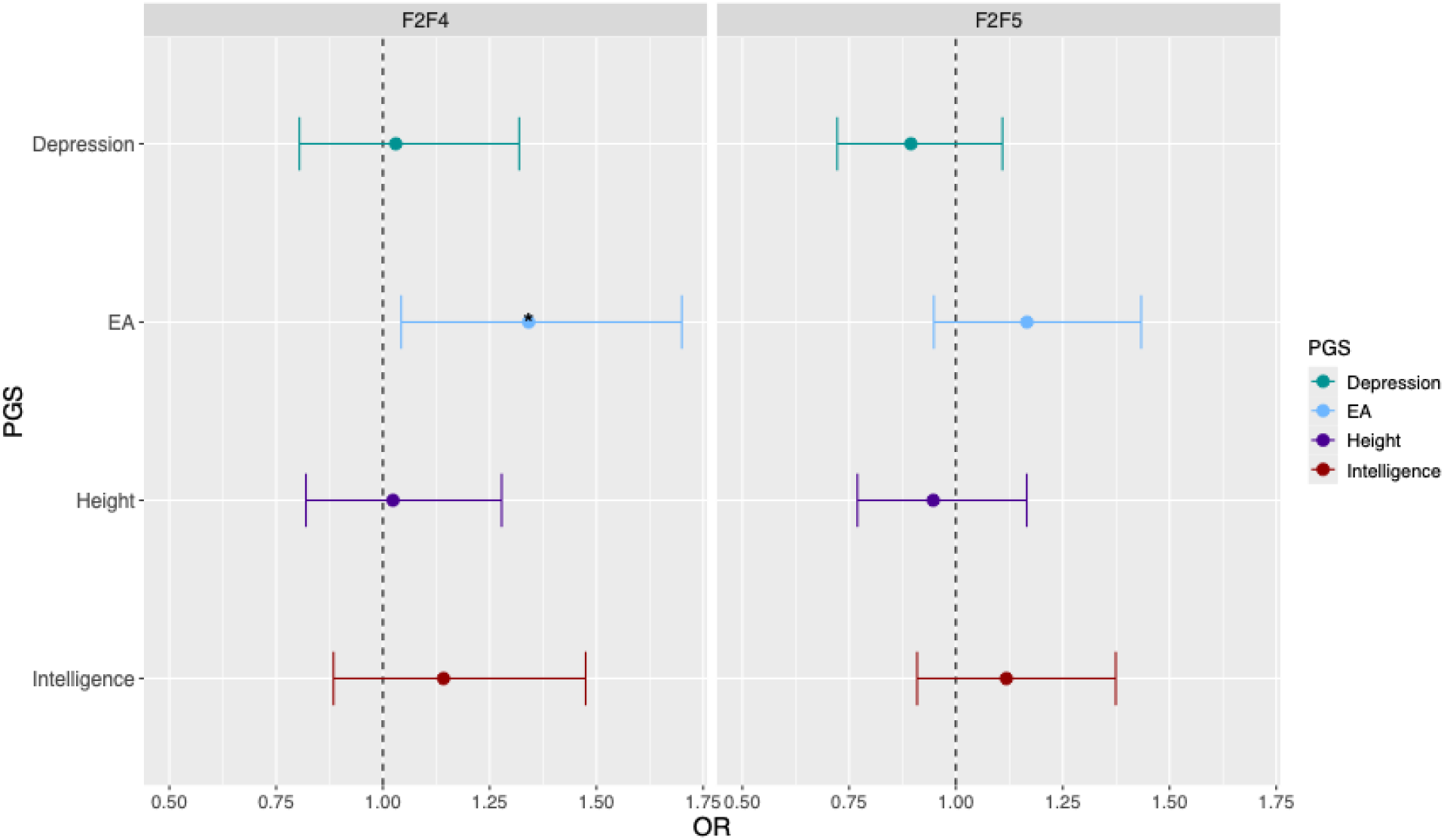
**Associations Between Polygenic Scores and Twins’ Study Participation in F2F4 and F2F5** *Note.* Analyses were conducted using logistic regression models adjusted for sex and birth cohort (cgr), with cluster-robust standard errors to account for family structure. The results shown here only include twins. See Supplementary Figure S2 for results including other family members. Asterisks indicate significance levels (* *p* < 0.05). EA = Educational attainment, OR = Odds ratio.

## Selective Sampling of TwinSNPs and TECS Subsamples is Reflected in Polygenic Scores

Furthermore, we compared four PGS (EA, intelligence, depression and height) for twins from the TECS sample (who are part of TwinSNPs) with the rest of the TwinSNPs sample. We ran linear models adjusting for sex, cohort and genetic PCs, effectively examining group differences between these two “subgroups”. TECS participants had significantly lower PGS for depression than the other TwinSNPs twins (*β* = –0.12, *SE* = 0.05, *p* = 0.015) and opposite trends were observed for the PGS for EA (*β* = 0.10, *SE* = 0.05, *p* = 0.060; Supplementary Figure S3). Sensitivity analyses, i.e., restricting to one random twin per family (for TECS vs TwinSNPs comparisons), restricting to DZ twins, or also including TwinSNPs participants that are not twins (siblings and parents), showed highly similar results (Supplementary Figures S4-S6).

## Integrating Genomic and DNA Methylation Data

Biological processes unfold across interconnected molecular layers, making the integration of multiple omics modalities essential for capturing the full complexity of human health and disease. Combing, for example, genotype and DNAm data can be used to enhance phenotype prediction or to assess the association between genetic and epigenetic factors, e.g., mQTL analysis as mentioned above. The examination of different omics layers in within-family designs is of further interest for causal inference or when studying direct and indirect genetic effects on DNAm (46,47). Some of these substantive research questions are currently ongoing in TwinLife. Rather than studying specific phenotypes, we here focus on initial technical results combining PGS and DNAm data in the form of epigenetic clocks and examine differences across timepoints (pre- and during the pandemic).

### Associations Between Polygenic Scores and Epigenetic Clocks

Whereas some studies have shown that PGS and MPS can capture independent components of phenotypes (48), DNAm is known to be influenced by genetics (49). Specifically, genetic effects on specific epigenetic clocks have been examined in a previous GWAS (21). As a proof of concept, we calculated three PGS for epigenetic clocks for which GWAS summary statistics are publicly available, i.e., for Hannum, Horvath, and GrimAge clocks (21). We tested associations between these three PGS and the calculated epigenetic clocks in the TECS sample using linear models for each time point (pre-pandemic and during the pandemic) to examine potential differences over time. In general, PGS for epigenetic clocks showed small associations with the target epigenetic clock after accounting for sex, age, cell-type composition and DNAm-based measure of smoking. At timepoint 1 (F2F3, pre-pandemic), none of the PGS was significantly associated with their respective epigenetic clock (see Supplementary Figure S9). At timepoint 2 (Cov3, during the pandemic), the PGS for GrimAge was significantly associated with the GrimAge and GrimAge2 clocks (*β* = 0.03, *SE* = 0.01, *p* = 0.010 and *β* = 0.04, *SE* = 0.01, *p* = 0.002, respectively; Figure 3). Results were highly similar when restricting to DZ twins (Supplementary Figures S10-S11), albeit showing larger standard errors due to the smaller sample sizes. This finding suggests that common genetic variants captured by PGS might be relatively more relevant for epigenetic changes during times when the total environmental variance is reduced due to universal environmental and societal shocks such as the COVID-19 pandemic.

**Figure 3.**
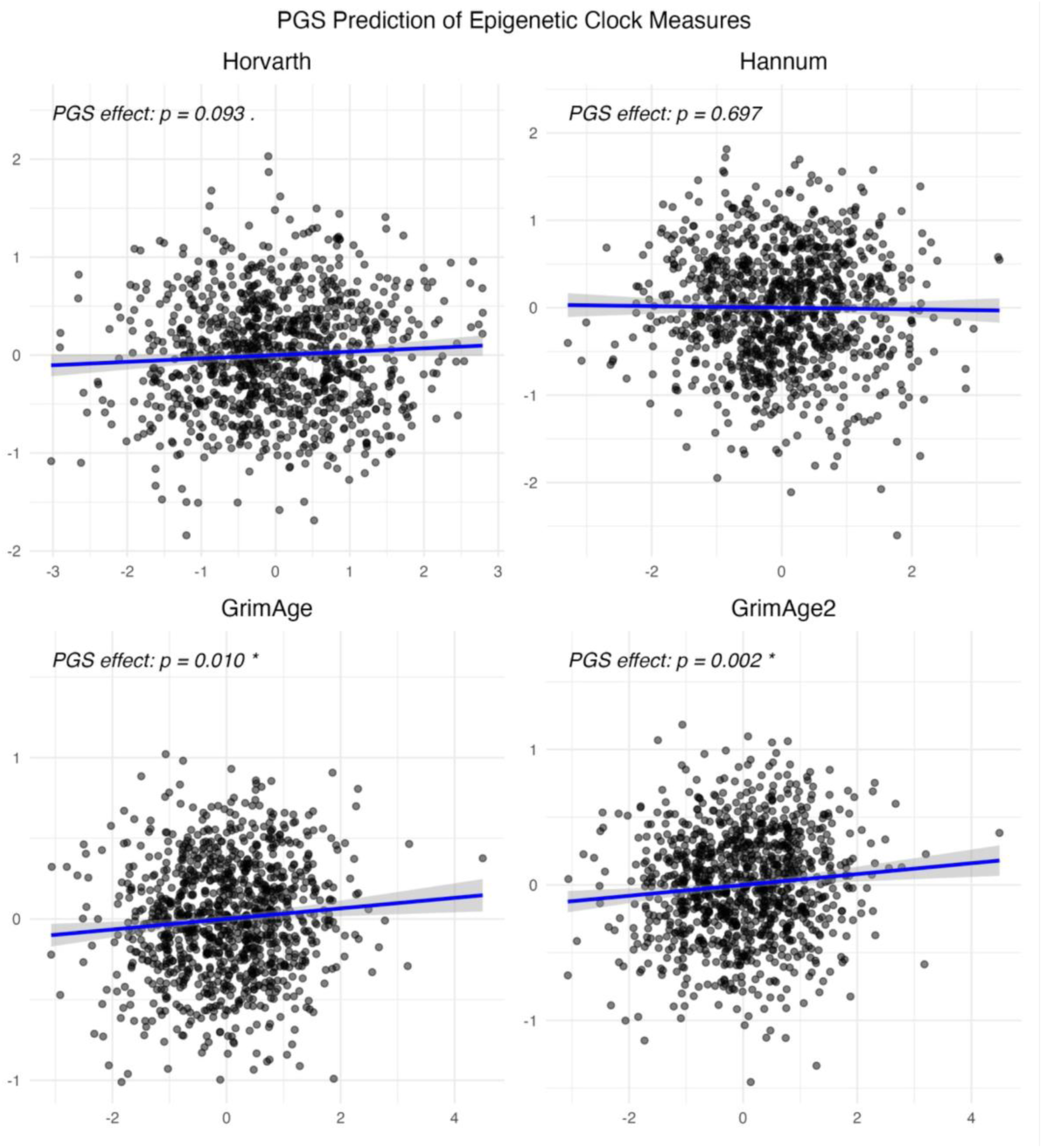
**Associations Between Polygenic Scores for Epigenetic Clocks and the Respective Epigenetic Clock in the TECS Sample at T2** *Note*. X-axes show the standardised PGS and y-axes show the respective standardised epigenetic clock.

## Strengths and Limitations

TwinSNPs and TECS provide unique opportunities to move beyond phenotypic associations and investigate underlying molecular pathways shaping development and socioeconomic disparities. Notably, the availability of genotype data in large extended twin samples is still rare. Using genetic data of TwinSNPs enables within-family genomic analyses such as examining direct and indirect genetic effects or intergenerational Mendelian randomisation (MR) to strengthen causal inference (50,51). In particular, causal inference methods using genotype data in twins can be applied, e.g., MR-DoC, combining advantages of both MR and relatedness information of twin samples (52,53). Genotype data can also be used to investigate gene-environment interactions both in unrelated individuals and within families. Furthermore, the TwinSNPs data can be integrated with DNAm data of TECS. This allows studying genetic and environmental contributions to DNAm using both molecular genetic approaches such as mQTL analysis as well as quantitative genetic approaches such as the twin family design. In addition, the prospective coverage of both pre- and post-COVID-19 periods in TECS provides the rare opportunity to examine the pandemic’s impact on DNAm, social inequality, and well-being. Combined with TwinLife’s longitudinal phenotypic data–spanning critical developmental periods and enriched with geo-referenced information–TwinSNPs and TECS thus provide a powerful platform for integrative analyses of genetic, epigenetic and environmental influences across development.

However, the molecular satellite data should also be considered in light of its limitations. First, both the TwinSNPs and TECS samples are selective, e.g., showing a higher prevalence of highly educated individuals in these subsamples (see also Table 1) which limits the investigation of adverse socioeconomic settings, in particular, in the context of the COVID-19 pandemic. However, this limitation could be addressed using sampling weights. In addition, DNAm data in TECS is only available for saliva, not for blood, which is the most extensively studied tissue in DNAm research and commonly used to train and validate MPS (54). Using blood has the benefit of analysing DNAm in different immunology-relevant cell types, either by isolating (55) or using computational modelling (56,57), but is harder to obtain and more invasive. Moreover, DNAm data currently is only generated for two timepoints which does not allow the investigation of long-term or delayed DNAm changes in relation to the pandemic.

## Outlook

In this Cohort Profile we have presented the data curation and initial analyses of the molecular genetic and epigenetic satellite projects TwinSNPs and TECS. There are several promising avenues that can build upon the saliva sample collection and DNA extraction in TwinLife. First, processing of additional saliva samples from F2F5 is ongoing and will further enlarge the genotyped TwinSNPs sample. Second, DNAm analysis of the third saliva sample (F2F5) is planned for the TECS sample, enabling longitudinal analysis of DNAm in a genetically informed design. Similarly, DNAm analysis for non-twin siblings is planned to enlarge the overall TECS sample. Third, generation of other omics data, e.g., whole-genome sequencing data from available saliva samples is possible and can deliver insights into the biological mechanisms of cognitive development and related phenotypes.

## Data Access

Currently, phenotype data are accessible free of charge via the GESIS data catalogue https://search.gesis.org/research_data/ZA6701 under protected access. Molecular genetic and epigenetic data can be used under collaboration and following an additional data transfer agreement. Pre-calculated PGS and epigenetic clocks can be shared with researchers applying for TwinLife data, as well as raw data. For enquiries, please contact. Further information can be found online https://www.twin-life.de/documentation/.

## Code Availability

Code for statistical analyses presented in this article can be found on GitHub https://github.com/AGForstner/TwinLife_CohortProfile.

## Ethics Approval

The TwinLife study was reviewed and approved by the German Psychological Society (Deutsche Gesellschaft für Psychologie; protocol number: RR 11.2009). The COVID-19 supplementary questionnaires were reviewed and approved by the Ethics Committee of the University of Bielefeld (Application No. 2020-106).

All participants were informed in writing about the aims of the TwinLife study, the incentives, and that their participation was voluntary and could be withdrawn at any time. In addition, in all face-to-face surveys, participants received a two-page leaflet summarizing the procedures concerning data protection and privacy and their rights to demand the deletion of all data collected in line with German data integrity laws. Written material was sent to participants before all interviews. At the beginning of each interview, informed verbal consent was obtained from participants (and the participants’ legal guardians when the participants were underage) and documented. Following the national legislation and the institutional requirements, written informed consent was not required to participate in this study. Nevertheless, for all supplementary online surveys, written informed consent was obtained from all participants.

The Ethics Committee of the Medical Faculty of the University of Bonn (No. 113/18) reviewed and approved the protocols for the molecular genetic analyses via saliva samples. Prior to the sampling, the participants received extensive written information on, e.g., the sampling procedures, the scope of the study, their right to refuse participation, and their right to withdraw their consent at any given time. For sampling molecular genetic data, written informed consent was obtained from all participants and the participant’s legal guardian (for minors).

## Author Contributions

CKLP, LF, AMS, CM and LW prepared the genetic and epigenetic data for TwinSNPs and TECS. MR prepared the phenotypic data. CKLP, LF and AMS performed the statistical analyses. CK, BM, FMS, MMN, EBB and AJF designed the projects, acquired funding, and supervised the core and satellite projects. CKLP, LF, AMS, AA, MD, MR, TR, and JI were involved in the implementation of the projects. EBB, DC and AJF supervised the current work. CKLP, LF and AMS drafted the manuscript and all authors critically reviewed and edited the manuscript.

## Supplementary Materials

Supplementary materials are available through the online version of this article.

## Acknowledgements

We thank Shirin Zare for performing the DNA extraction and Dr. Friederike David for sharing code for PGS calculation. We also thank Angelina Vogelsang for supporting the DNA methylation profiling. We gratefully acknowledge the access to the bonna cluster at the University of Bonn and the support provided by the HPC@HRZ team at the University of Bonn. We are also thankful for all the other contributions of researchers and student assistants that were involved in the implementation of the projects.

