## Supplementary Materials for "(Epi-)Genomic Data in the German TwinLife Study: TwinSNPs and TECS Cohort Profiles"

### Supplementary Note

#### *Genotyping and Imputation*

DNA was extracted using standard protocols. Genotyping was performed using Global Screening Arrays (GSA+MD-24v3.0-Psych-24v1.1, Illumina, San Diego, CA, USA), with additional content relevant to psychiatric disorders. As also described previously (Deppe et al., 2026), quality control (QC) of genome-wide genotype data was performed using PLINK (Purcell et al., 2007) and R (v4.0.3; R Core Team, 2024). Raw genotype data were available for 5,927 samples. After filtering, this resulted in 5,861 individuals based on a high overall genotyping rate of 98% and autosomal heterozygosity deviation (FHET) within +/- 0.20. Variants with a call rate < 98%, variants deviating from Hardy-Weinberg equilibrium with a p-value $< 1 \times 10^{-6}$ among unrelated individuals (considering one sample per family selected according to the highest genotyping rate) or a minor allele frequency (MAF) < 0.001 were removed. Post-QC data were phased using Eagle (v2.4.1; Loh et al., 2016) and imputed using Minimac (v4; Das et al., 2016). 1000 Genomes Phase 3 data (1000Gv3; $N$ = 2,504, Fairley et al., 2020) was used as imputation reference panel after excluding ambiguous variants (A/T, C/G) to avoid potential strand issues alignment between the reference and target data. Without filtering based on imputation quality (INFO) scores, the imputed data set consisted of 11,108,941 variants.

#### *Further Sample-Level Filtering for PGS Analyses Based on Sex, Relatedness and Population Substructure Analyses*

Concordance of genetically inferred sex and self-reported sex was checked using PLINK v1.9. Samples with undetermined genetically inferred sex and samples with a sex mismatch were excluded.

As the TwinSNPs sample consists of twins and their family members, a relatedness check was included in the QC. For this purpose, KING (Manichaikul et al., 2010) was used to compute the kinship coefficients between samples and infer familial relationships from genotype data. Samples with a discordant genetically inferred family structure (reflected by pairwise KING relatedness coefficients and KING-generated family IDs) with respect to reported relatedness (reflected by the fid and ptyp SUF variables) were excluded, with the exception of zygosity mismatches. We retained monozygotic pairs with an inferred genetic zygosity discordant with the reported zygosity if the genetically inferred family structure was otherwise consistent with the self-report. In these cases, the zygosity variable was corrected to include the genetically inferred zygosity.

To partially adjust for population stratification in subsequent analyses, the first 10 genetic principal components (PCs) were derived by projecting the samples into the PC space defined using 1000Gv3 as reference panel. Each TwinSNPs sample was assigned to the closest superpopulation and subpopulation of the 1000Gv3 reference data based on the smallest Euclidean distance to the corresponding population centroids in the first 10 PCs space (or not assigned if the shortest distance exceeds the threshold derived from the distribution of the overall population centroid distances, corresponding to a fixation index of .02). In addition, the principal component analysis was used to flag PC outliers, defined as samples whose PC values deviated more than 6 standard deviations from the mean of the first 10 PCs.

***Polygenic Score Calculation***

Preprocessing of GWAS summary statistics comprised filtering the genetic variants for (i) an imputation info score ≥ 0.6 if available, (ii) a minor allele frequency ≥ 0.01 in the 1000Gv3 European LD reference panel, and (iii) presence in both the summary statistics and the imputed genotype data of our cohort. For the remaining variants, weights were then estimated based on their GWAS effect sizes using PRS-CS (Ge et al., 2019), with the 1000Gv3 European LD reference panel and default parameters (PRS-CS auto). PRS-CS uses Bayesian regression that applies continuous shrinkage (CS) priors to estimate SNP effects. The auto model learns the phi parameter from the input summary statistics while adjusting for LD using an external reference panel. In case of a negative association between the effect allele and phenotype, effect and non-effect alleles were flipped so that posterior effect sizes were always positive. From the estimated weights and the imputed genotype data, polygenic scores (PGS) were calculated using plink v1.9 (--score). Where GWAS data was not available for optimisation based on the PRS-CS pipeline, we used PGS score weight files (if available) to calculate the PGS directly using the plink --score function for our target dataset. The score weights are usually estimated based on GWAS summary statistics (which contain effect sizes and standard errors of each tested SNP) in an independent target sample.

***Performance of Epigenetic Clocks***

In addition to the correlation with chronological age described in the main text, we provide the median absolute error (MAE, i.e., absolute median difference between chronological and epigenetic age) and median deviation (i.e., median of chronological and epigenetic age) separately for each cohort. The MAE and median deviation ($\delta$) can indicate potential cohort-specific biases in the age estimation. As illustrated in Supplementary Figures S6–S7, Horvath's clock, e.g., performed well in all age groups (children: MAE = 2.9 to 2.8, delta = 3.1 to 2.9; adolescents: MAE = 3.3 to 3.7, delta = 2.2 to 2.7; adults: MAE = 3.2 to 3.5, delta = 0.8 to 1.4) whereas PedBE performed well in children and adolescents but not in adults (children: MAE = 1.1 to 2.0, $\delta$ = 1.0 to 2.1; adolescents: MAE = 2.3 to 2.9, $\delta$ = –2.9 to –2.2; adults: MAE = 8.2 to 9.5, $\delta$ = –9.9 to –8.7). The latter is consistent with the specific purpose of age prediction in children that PedBE was designed for.

### Supplementary Tables

**Supplementary Table S1**

*Overview of Currently Available Methylation Profile Scores (MPS) for TECS*

| Category | MPS (ref) | Description | Available CpGs (%) | Stability over time | Correlation with respective phenotype | |
|---|---|---|---|---|---|---|
| | | | | ICC[1] | T1 | T2 |
| Educational & cognitive traits | Epigenetic-g (McCartney et al., 2022) | DNAm surrogate for general cognitive ability. It was computed as the weighted sum of betas values associated with a factor for general cognitive ability in McCartney et al. using the code provided at https://github.com/marioni-group/ewas_of_cognitive_function. A higher score indicates DNAm changes associated with a higher cognitive ability. | 99.98% | 0.73 | | |
| Physical traits | BMI (Do et al., 2023) | DNAm surrogate for BMI. It was computed as the weighted sum of beta values derived from the EWAS summary statistics by Do et al. A higher score indicates DNAm changes associated with a higher BMI. | 92.95% | 0.55 | 0.1 | NA |
| | Puberty (deSteiguer et al., 2025) | | 100% | 0.89 | See details in deSteiguer et al. (2025) | |
| Protein biomarkers | CRP (Barker et al., 2018) | DNAm surrogate for CRP. It was computed as the weighted sum of beta values derived from the EWAS summary statistics by Barker et al. A higher score indicates DNAm changes associated with higher levels of circulating CRP. | 83.33% | 0.44 | NA | NA |

| | | | | | | |
|---|---|---|---|---|---|---|
| | CRP (Ligthart et al., 2016) | DNAm surrogate for CRP. It was computed as the weighted sum of beta values derived from the EWAS summary statistics by Lighart et al. A higher score indicates DNAm changes associated with higher levels of circulating CRP. | 94.95% | 0.32 | NA | NA |
| Lifestyle exposure | Smoking CpG (Philibert et al., 2013) | Beta value of the individual CpG site cg05575921. De-methylation of the probe cg05575921 has been shown to be sensitive to smoking frequency in both saliva and blood. Values were reverse-coded so that a higher beta value of this probe indicates higher smoking exposure. | 100% | 0.46 | NA | NA |
| Aging: biological markers | Telomere length (Lu, Seeboth, et al., 2019) | DNAm surrogate for telomere length. It was computed as the weighted sum of beta values linked to telomere length in Lu et al. using the R-package methylclock v1.01 (Pelegí-Sisó et al., 2021). A higher score indicates DNAm changes associated with longer terminal restriction fragments (in kilobases). | 100% | 0.49 | NA | NA |
| | GrimAge (Lu, Quach, et al., 2019) | Epigenetic age estimate. It was computed as the sum of several DNAm surrogate scores for protein biomarkers (e.g., leptin, cholesterol), female, and smoking based on code obtained by personal correspondence with Lu et al. A higher score indicates a higher epigenetic age (in years). | 99.90% | 0.82 | 0.9 | 0.85 |
| | PC GrimAge (Higgins-Chen et al., 2022) | Epigenetic age estimates derived from principal components instead of individual GrimAge beta values. It was computed using the code provided at https://github.com/HigginsChenLab/PC-Clocks. A higher score indicates a higher epigenetic age. | 100 % | 0.92 | 0.92 | 0.89 |

| | | | | | | |
|---|---|---|---|---|---|---|
| | GrimAge2 (Lu et al., 2022) | Epigenetic age estimate. It was computed as the sum of several DNAm surrogate scores for protein biomarkers (e.g., leptin, cholesterol), sex, and smoking based on code obtained by personal correspondence with Lu et al. It includes two additional protein markers (hemoglobin A1C, CRP) compared to GrimAge. A higher score indicates a higher epigenetic age (in years). | 99.0% | 0.71 | 0.8 | 0.72 |
| | PhenoAge (Levine et al., 2018) | Epigenetic age estimate. It was computed as the weighted sum of the DNAm surrogates for 9 blood biomarkers (e.g., lymphocyte count, CRP) using the R-package methylclock v1.01. A higher score indicates a higher epigenetic age (in years). | 100% | 0.8 | 0.83 | 0.81 |
| | PC phenoAge (Higgins-Chen et al., 2022) | Epigenetic age (in years) derived from principal components instead of individual PhenoAge beta values. It was computed using the code provided at https://github.com/HigginsChenLab/PC-Clocks. A higher score indicates a higher epigenetic age. | 100 % | 0.79 | 0.83 | 0.75 |
| | DunedinPACE (Belsky et al., 2022) | Epigenetic pace of aging where a value of 1 represents the same rate of aging as 1 chronological year. It was computed as the weighted sum of DNAm surrogate markers for longitudinal biomarker trajectories (e.g., CRP, BMI) using the code provided at https://github.com/danbelsky/DunedinPACE.git. | 100% | 0.36 | NA | NA |
| Aging: chronological age | Horvath DNAmAge (Horvath, 2013) | Epigenetic age estimate. It was computed as the weighted sum of beta values associated with chronological age in Horvath et al. using the R-package methylclock v1.01. A higher score indicates a higher epigenetic age (in years). | 94.6% | 0.88 | 0.87 | 0.87 |

| | | | | | | |
|---|---|---|---|---|---|---|
| | PC Horvath (Higgins-Chen et al., 2022) | Epigenetic age estimates derived from principal components instead of individual Horvath beta values. It was computed using the code provided at https://github.com/HigginsChenLab/PC-Clocks. A higher score indicates a higher epigenetic age. | 100% | 0.84 | 0.84 | 0.83 |
| | SkinHorvath (Horvath et al., 2018) | Epigenetic age estimate. It was computed as the weighted sum of beta values associated with chronological age in Horvath et al. using the R-package methylclock v1.01. A higher score indicates a higher epigenetic age (in years). | 100% | 0.94 | 0.93 | 0.94 |
| | PC SkinHorvath (Higgins-Chen et al., 2022) | Epigenetic age estimates derived from principal components instead of individual SkinHorvath beta values. It was computed using the code provided at https://github.com/HigginsChenLab/PC-Clocks. A higher score indicates a higher epigenetic age. | 100% | 0.85 | 0.83 | 0.83 |
| | BLUP (Zhang et al., 2019) | Epigenetic age estimate. It was computed as the weighted sum of beta values associated with chronological age in Zhang et al. using the code provided at https://github.com/qzhang314/DNAm-based-age-predictor. A higher score indicates a higher epigenetic age (in years). | 99.90% | 0.95 | 0.93 | 0.94 |
| | EN (Zhang et al., 2019) | Epigenetic age estimate. It was computed as the weighted sum of beta values associated with chronological age in Zhang et al. using the code provided at https://github.com/qzhang314/DNAm-based-age-predictor. A higher score indicates a higher epigenetic age (in years). | 100% | 0.95 | 0.94 | 0.95 |

| | | | | | | |
|---|---|---|---|---|---|---|
| | PedBE (McEwen et al., 2020) | Epigenetic age estimate. It was computed as the weighted sum of beta values associated with chronological age in McEwen et al. using the R-package methylclock v1.01. A higher score indicates a higher epigenetic age (in years). | 100% | 0.83 | 0.86 | 0.81 |
| | Wu (Wu et al., 2019) | Epigenetic age estimate (in years). It was computed as the weighted sum of beta values associated with chronological age in Wu et al. using the R-package methylclock v1.01. A higher score indicates a higher epigenetic age (in years). | 96.40% | 0.72 | 0.78 | 0.71 |
| | Hannum (Hannum et al., 2013) | Epigenetic age estimate. It was computed as the weighted sum of beta values associated with chronological age in Hannum et al. using the R-package methylclock v1.01. A higher score indicates a higher epigenetic age (in years). | 91.54% | 0.85 | 0.85 | 0.84 |
| | PC Hannum (Higgins-Chen et al., 2022) | Epigenetic age estimates derived from principal components instead of individual Hannum beta values. It was computed using the code provided at https://github.com/HigginsChenLab/PC-Clocks. A higher score indicates a higher epigenetic age. | 100% | 0.89 | 0.89 | 0.85 |
| | | | | | % correctly identified | |
| Other | Zygosity (van Dongen et al., 2021) | DNAm surrogate for zygosity, computed based on the EpiPredictorMZtwin.R script by van Dongen et al. | 100% | 0.90 | MZ: 82.26<br>DZ: 50.1 | MZ: 83.33<br>DZ: 44.38 |

*Note.* [1]The intra-class coefficient was computed across all participants with two DNAm samples (n = 1,047) using the icc function of the R-package *irr*, set to two-way model, consistency, and single unit to estimate the stability of the first measurement over time. CRP = C-reactive protein, BMI = body-mass-index, PC = principal component, EN = elastic net, BLUP = best linear unbiased prediction, PedBE = Pediatric-Buccal-Epigenetic, MZ = monozygotic, DZ = dizygotic, NA = not available.

### Supplementary Figures

#### Supplementary Figure S1

*Extended Twin Family Design in TwinLife*

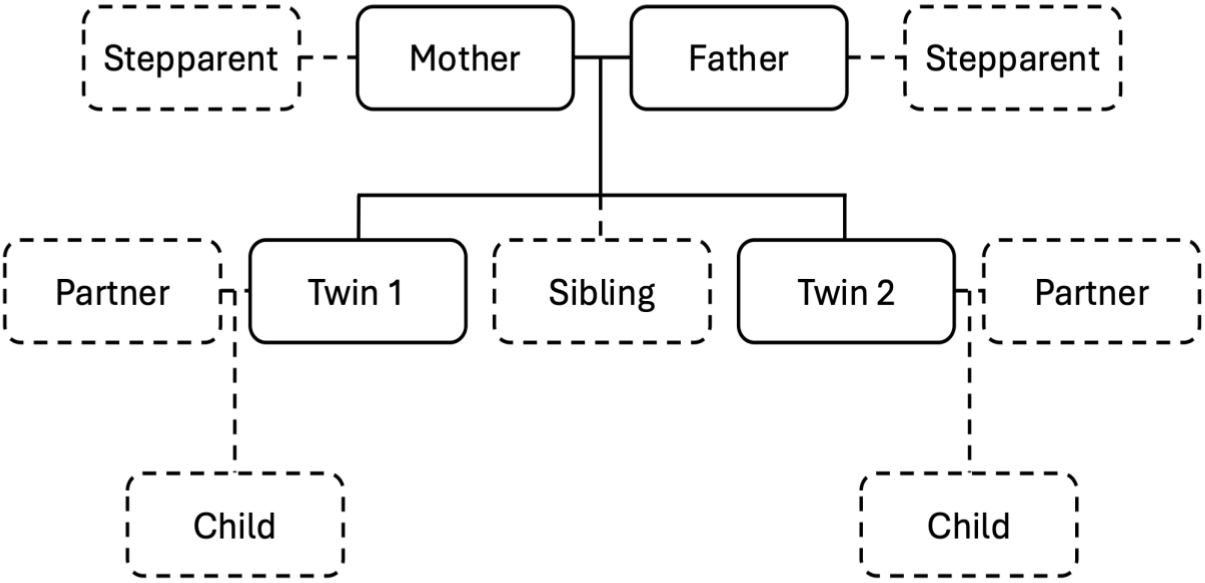


#### Supplementary Figure S2

*Associations Between Polygenic Scores and Participation of all Individuals*

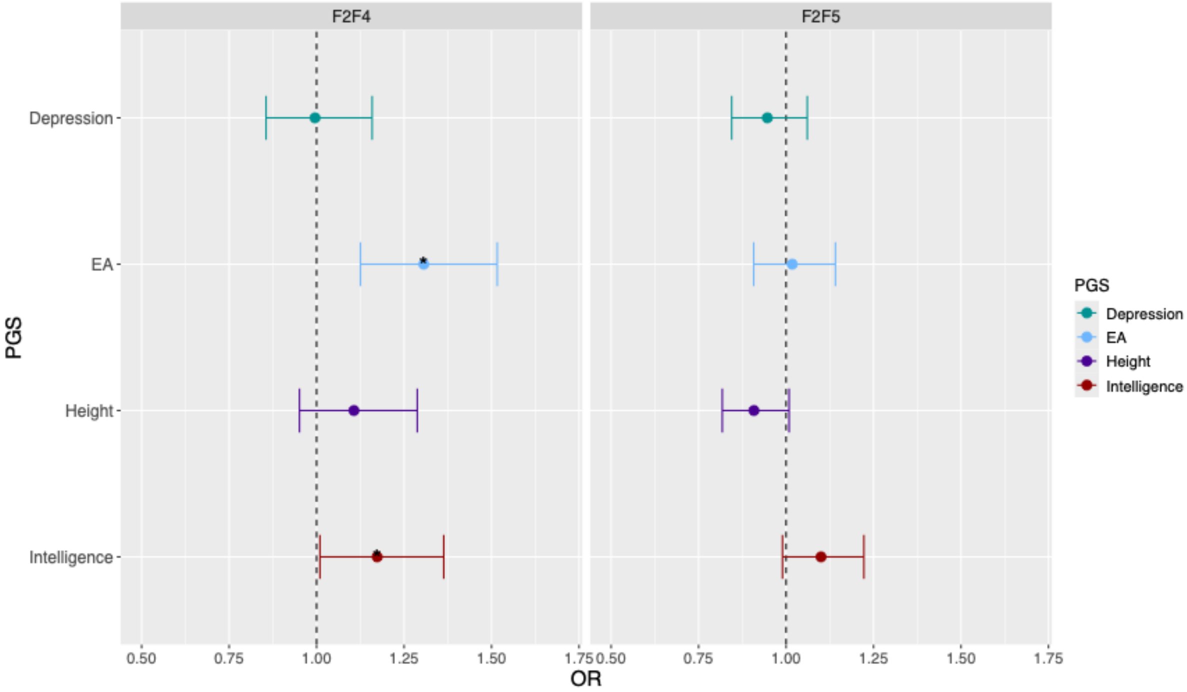

**Supplementary Figure S3**

*Differences in Polygenic Scores Between TECS and Other TwinSNPs Participants*

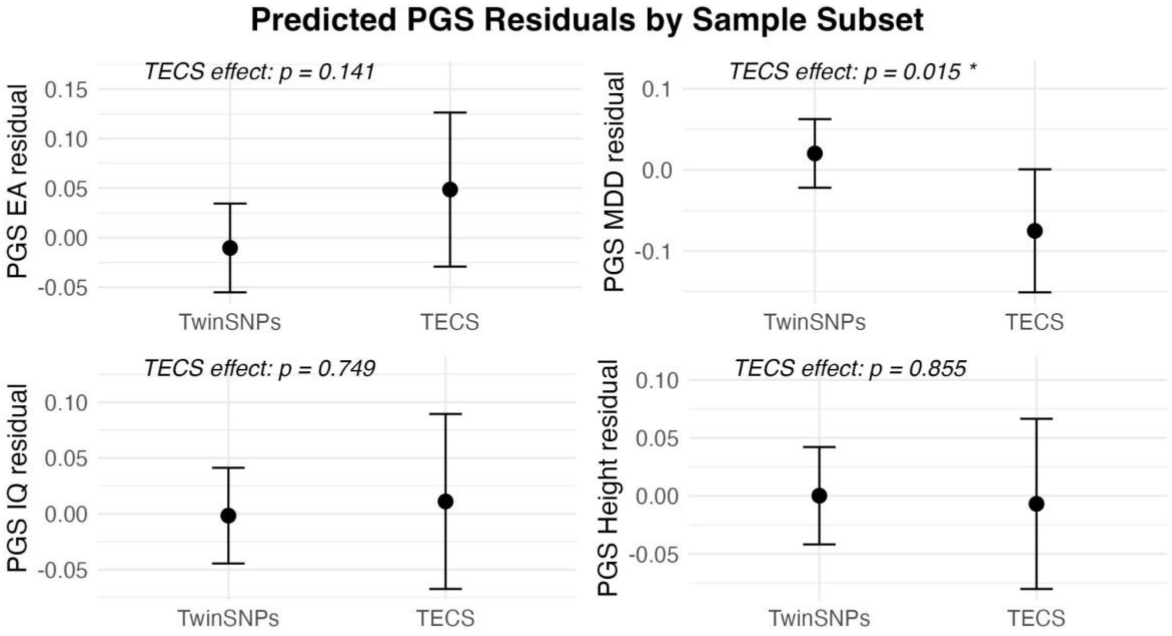


*Note.* Predicted polygenic score (PGS) residuals are shown for four traits (EA = Educational Attainment, MDD = Major Depressive Disorder, IQ = Intelligence, and Height), comparing participants from the TwinSNPs and TECS subsamples. Analyses were conducted using linear regression models adjusted for sex and birth cohort (cgr), with standard errors computed using cluster-robust variance estimation to account for family structure. The "TECS effect" p-values represent results from a cluster-robust t-test on the regression coefficient for the TECS sample indicator, testing whether PGS residuals differ significantly between TECS and TwinSNPs individuals. Asterisks indicate significance levels (* $p < 0.05$).

**Supplementary Figure S4**

*Differences in Polygenic Scores Between TECS Twins and TwinSNPs Twins and Non-Twin Siblings*

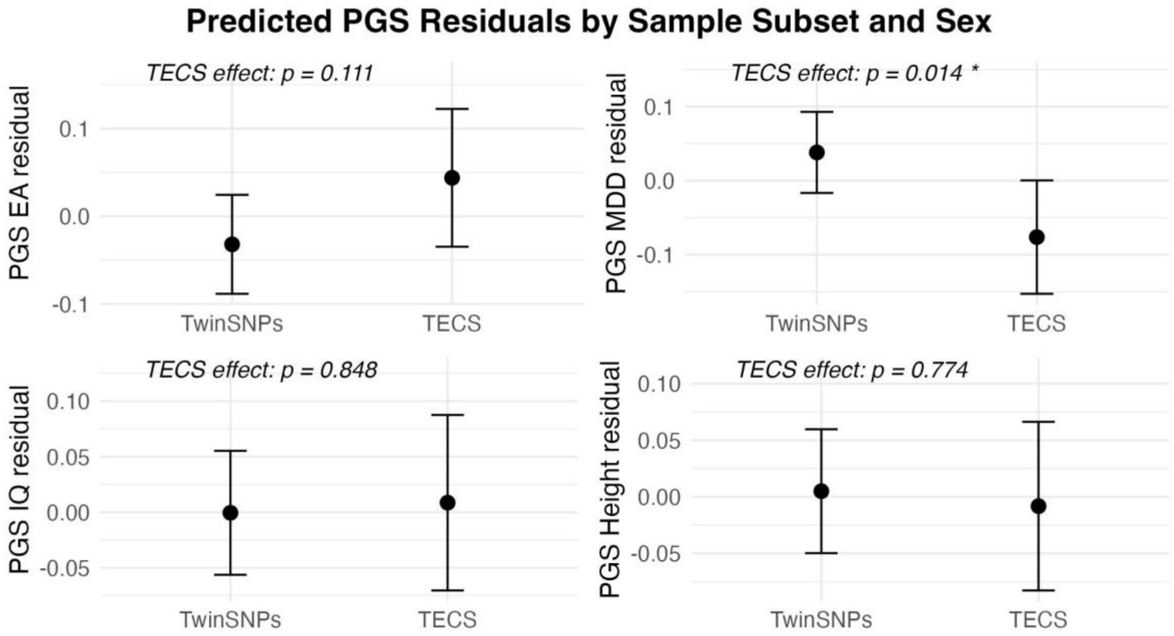


**Supplementary Figure S5**

*Differences in Polygenic Scores Between TECS and TwinSNPs MZ and DZ Twins*

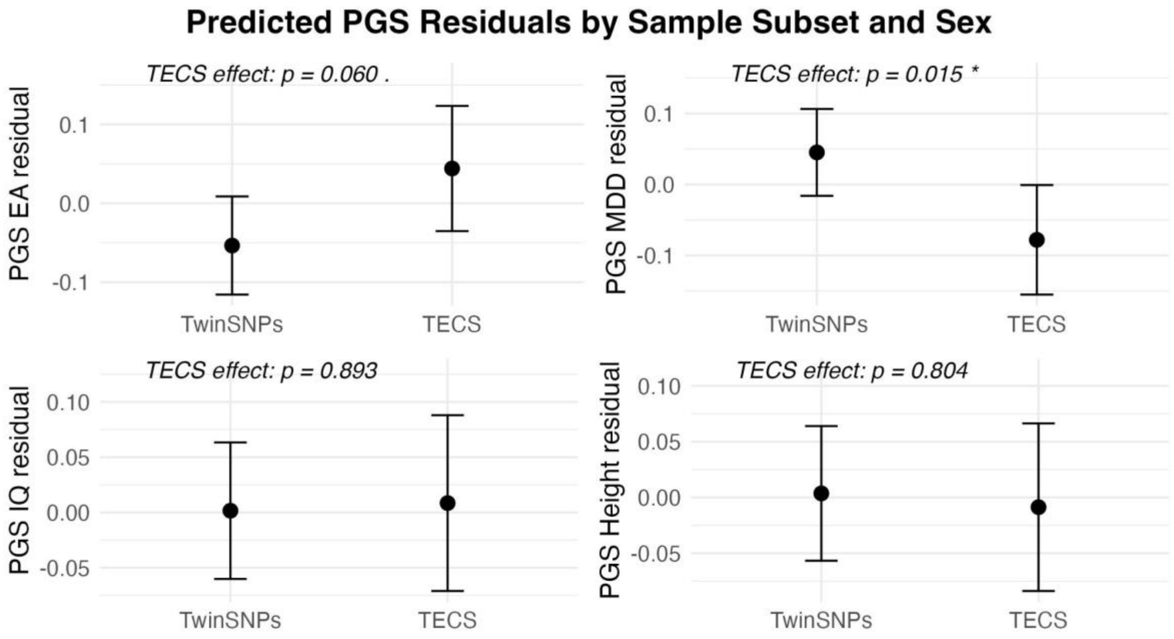

**Supplementary Figure S6**

*Differences in Polygenic Scores Between TECS and TwinSNPs DZ Twins*

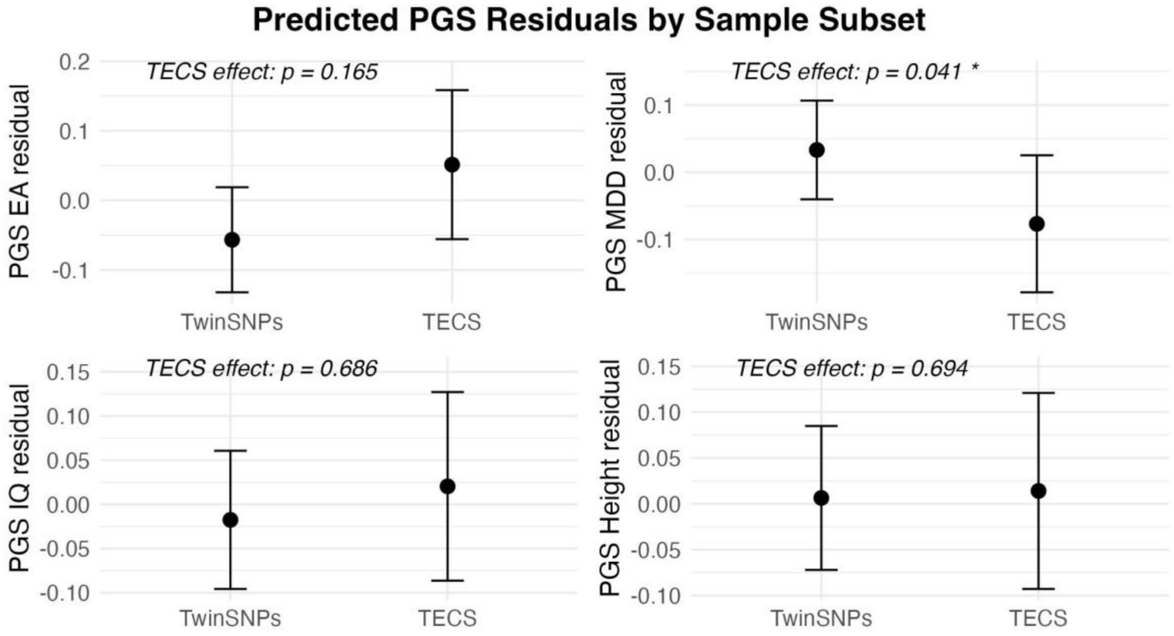

**Supplementary Figure S7**

*Median Absolute Error by Age Group*

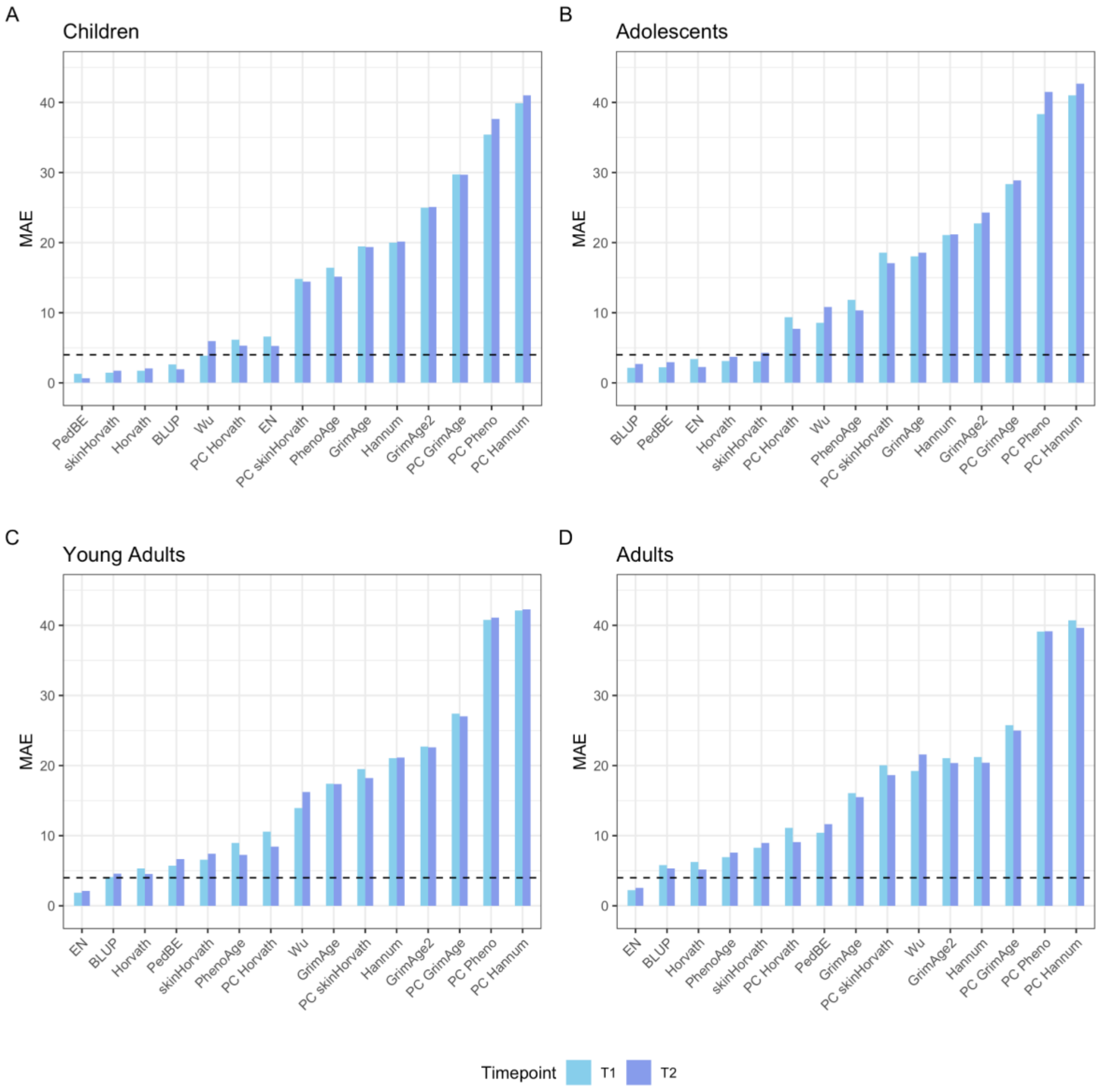


*Note.* Median absolute errors (absolute difference between chronological and epigenetic age) are presented for (A) children, (B) adolescents, and (C) adults at baseline and follow-up.

**Supplementary Figure S8**

*Mean Deviation by Age Group*

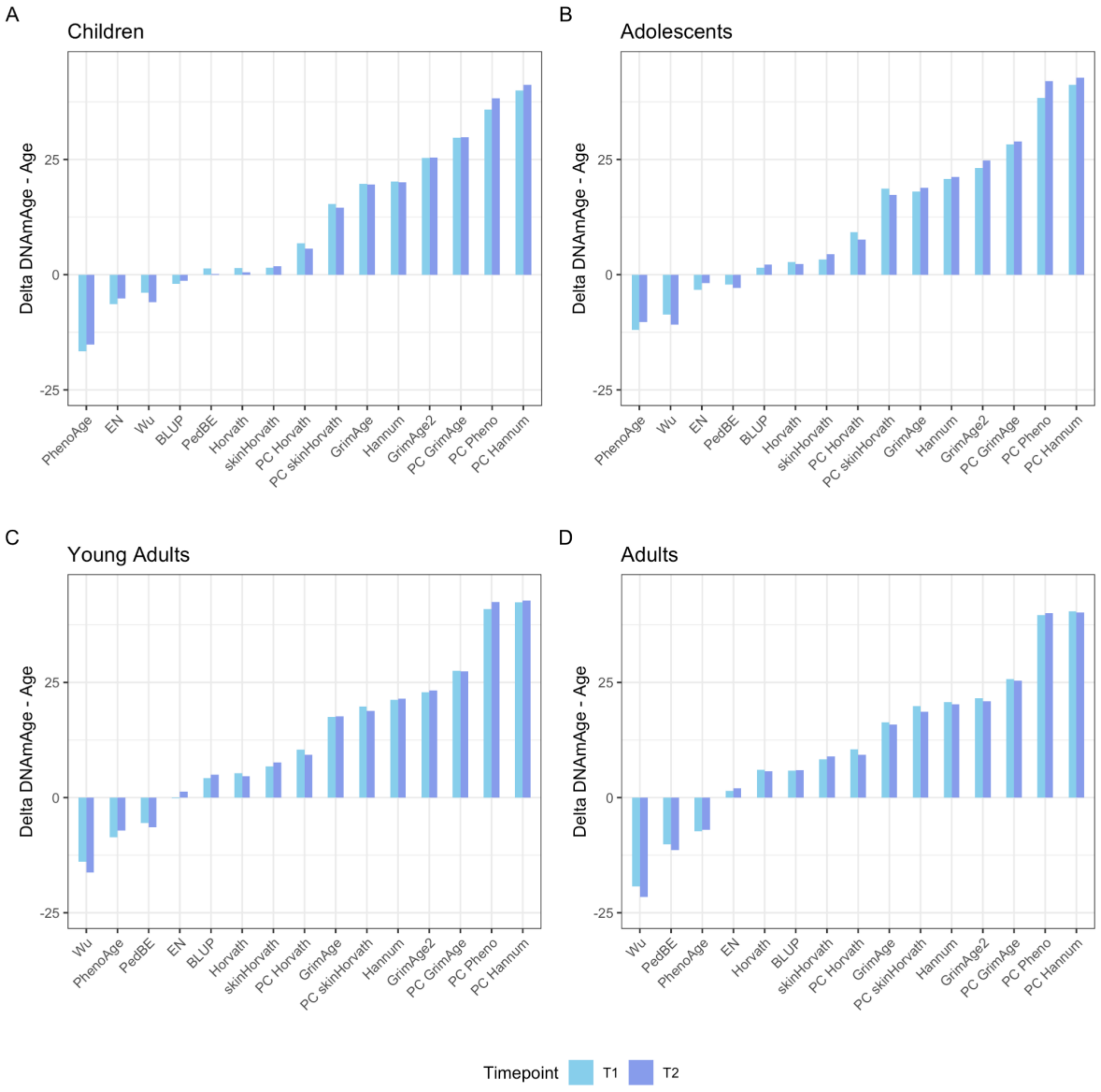


*Note.* Mean deviation (epigenetic minus chronological age) are presented for (A) children, (B) adolescents, and (C) adults at baseline and follow-up.

**Supplementary Figure S9**

*Associations Between Polygenic Scores for Epigenetic Clocks and the Respective Epigenetic Clock in the TECS Sample at T1*

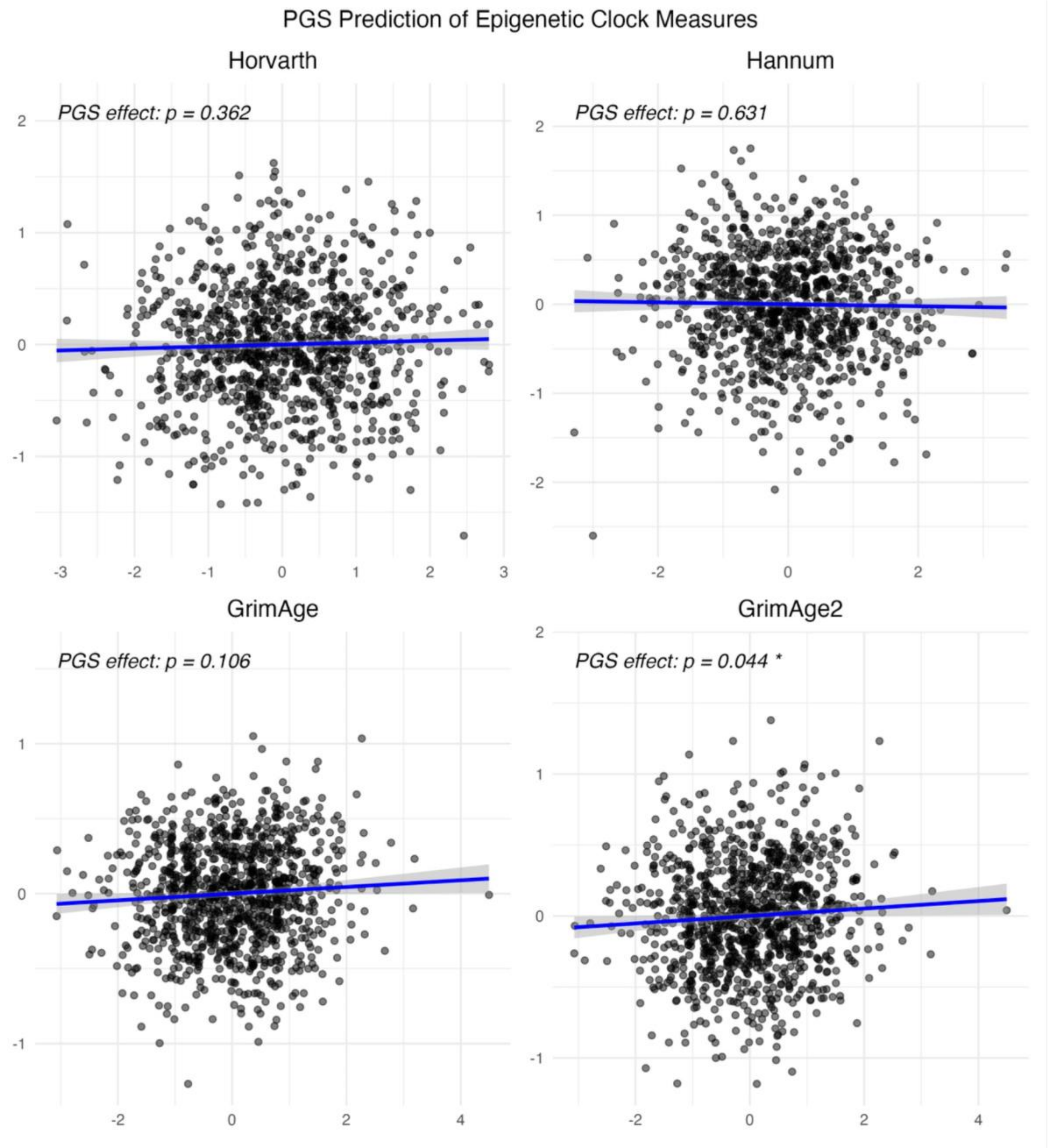

**Supplementary Figure S10**

*Associations Between Polygenic Scores for Epigenetic Clocks and the Respective Epigenetic Clock in the TECS Sample at T1, DZs only*

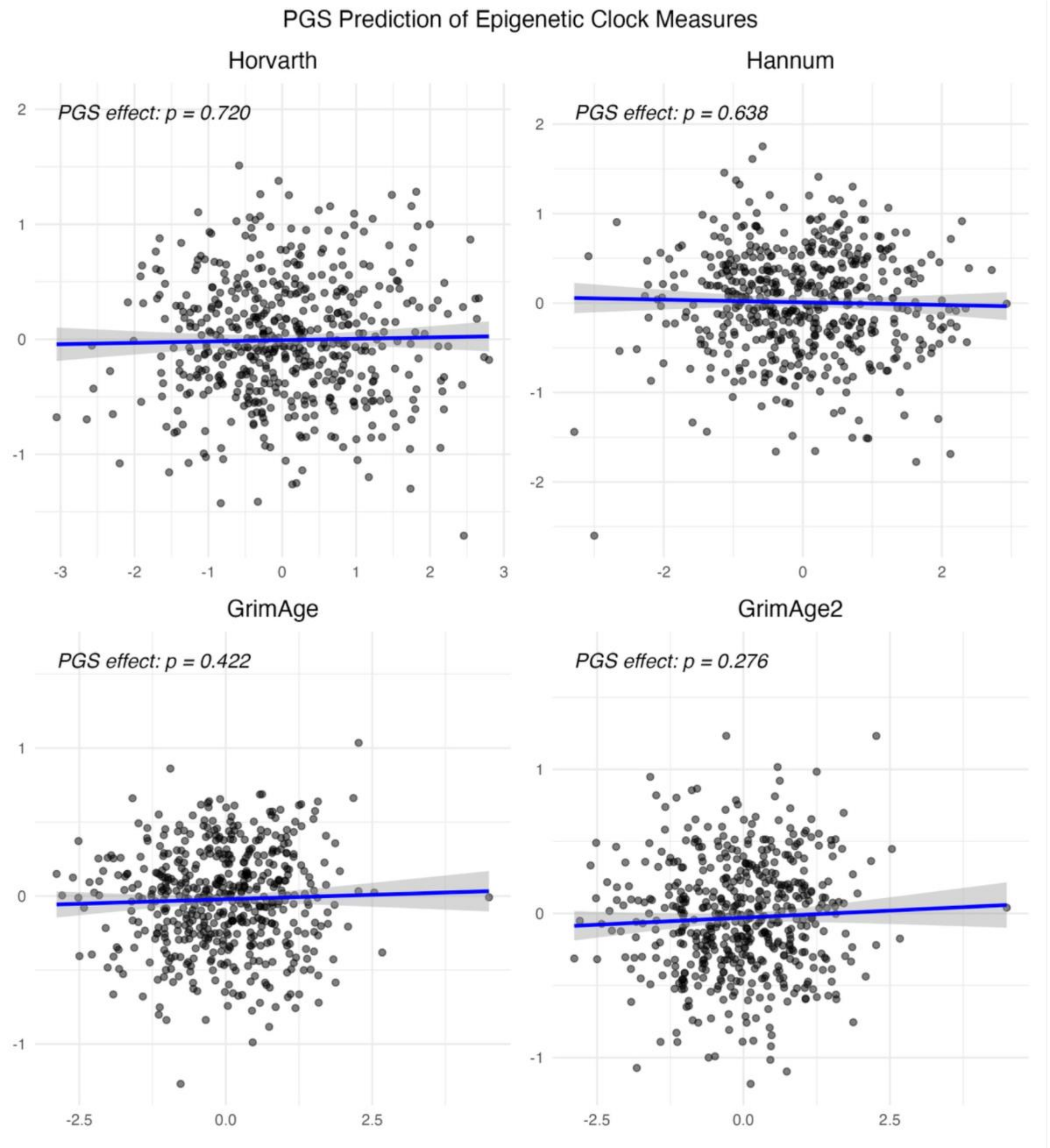

**Supplementary Figure S11**

*Associations Between Polygenic Scores for Epigenetic Clocks and the Respective Epigenetic Clock in the TECS Sample at T2, DZs only*

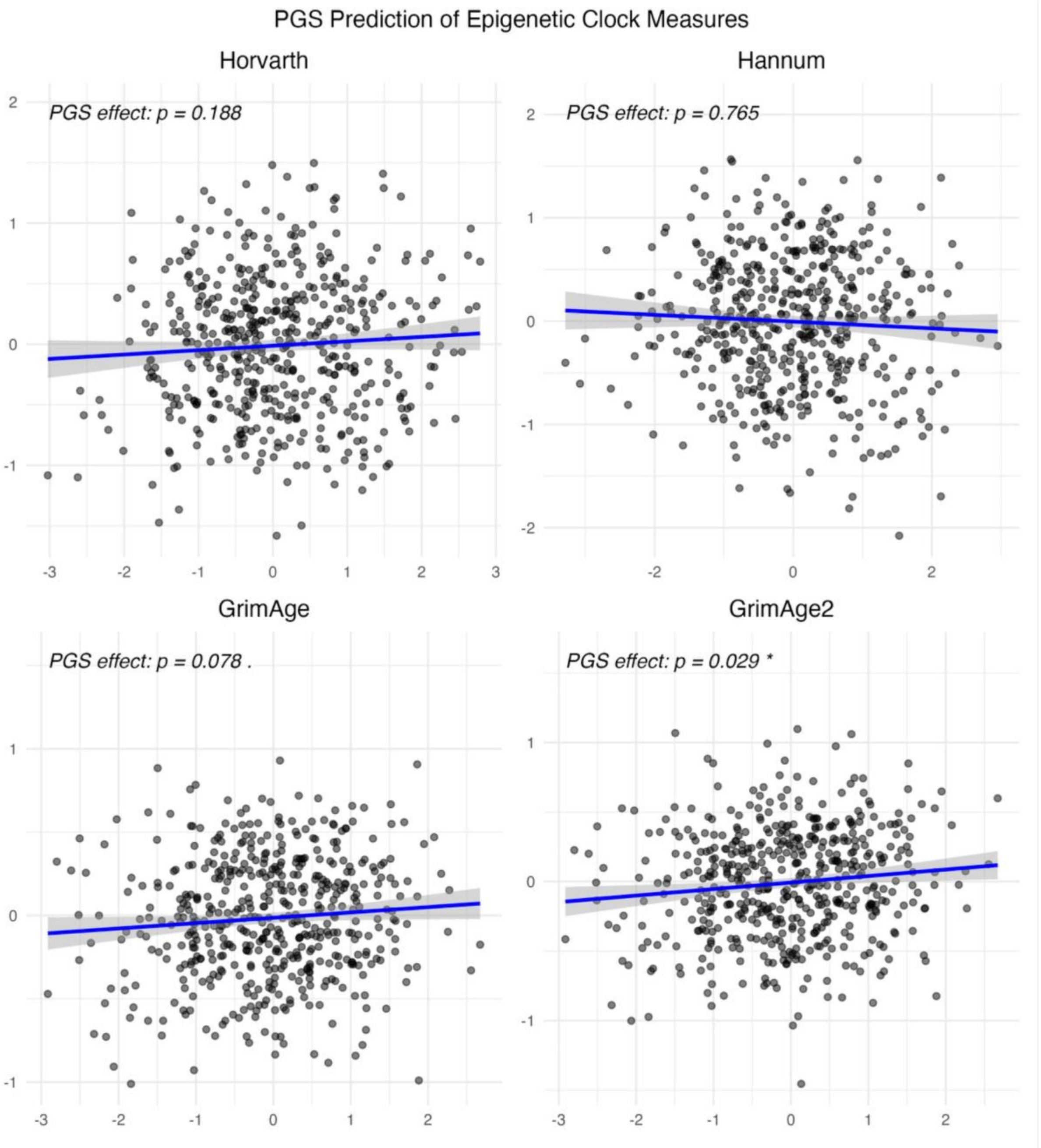
